# WNT2B Deficiency Causes Increased Susceptibility to Colitis in Mice and Impairs Intestinal Epithelial Development in Humans

**DOI:** 10.1101/2023.04.21.537894

**Authors:** Amy E. O’Connell, Sathuwarman Raveenthiraraj, Comfort Adegboye, Wanshu Qi, Radhika S. Khetani, Akaljot Singh, Nambirajam Sundaram, Chidera Emeonye, Jasmine Lin, Jeffrey D. Goldsmith, Jay R. Thiagarajah, Diana L. Carlone, Jerrold R. Turner, Pankaj B. Agrawal, Michael Helmrath, David T. Breault

**Affiliations:** Division of Newborn Medicine, Boston Children’s Hospital, Boston MA; Division of Endocrinology, Boston Children’s Hospital, Boston MA; Division of Genetics and Genomics, Boston Children’s Hospital, Boston MA; Department of Pathology, Boston Children’s Hospital, Boston MA; Division of Gastroenterology, Boston Children’s Hospital, Boston MA; Harvard T.H. Chan School of Public Health, General, and Thoracic Surgery, Cincinnati Children’s Hospital, Cincinnati, OH; Department of Pediatric, General, and Thoracic Surgery, Cincinnati Children’s Hospital, Cincinnati, OH; Center for Stem Cell and Organoid Medicine, Cincinnati Children’s Hospital, Cincinnati, OH; Laboratory of Mucosal Barrier Pathobiology, Department of Pathology and Medicine, Brigham & Women’s Hospital and Harvard Medical School; Department of Pediatrics, Harvard Medical School, Boston, MA; The Manton Center for Orphan Disease Research at Boston Children’s Hospital, Boston, MA; Division of Neonatology, Department of Pediatrics, University of Miami Miller School of Medicine and Holtz Children’s Hospital, Jackson Health System, Miami, FL; Harvard Stem Cell Institute, Cambridge, MA

**Keywords:** colon, colitis, WNT2B, WNT3, intestinal stem cells, intestinal development

## Abstract

**Background and aims:** WNT2B is a canonical Wnt ligand previously thought to be fully redundant with other Wnts in the intestinal epithelium. However, humans with WNT2B deficiency have severe intestinal disease, highlighting a critical role for WNT2B. We sought to understand how WNT2B contributes to intestinal homeostasis.

**Methods:** We investigated the intestinal health of *Wnt2b* knock out (KO) mice. We assessed the impact of inflammatory challenge to the small intestine, using anti-CD3ε antibody, and to the colon, using dextran sodium sulfate (DSS). In addition, we generated human intestinal organoids (HIOs) from WNT2B-deficient human iPSCs for transcriptional and histological analyses.

**Results:** Mice with WNT2B deficiency had significantly decreased *Lgr5* expression in the small intestine and profoundly decreased expression in the colon, but normal baseline histology. The small intestinal response to anti-CD3ε antibody was similar in *Wnt2b* KO and wild type (WT) mice. In contrast, the colonic response to DSS in *Wnt2b* KO mice showed an accelerated rate of injury, featuring earlier immune cell infiltration and loss of differentiated epithelium compared to WT. WNT2B-deficient HIOs showed abnormal epithelial organization and an increased mesenchymal gene signature.

**Conclusion:** WNT2B contributes to maintenance of the intestinal stem cell pool in mice and humans. WNT2B deficient mice, which do not have a developmental phenotype, show increased susceptibility to colonic injury but not small intestinal injury, potentially due to a higher reliance on WNT2B in the colon compared to the small intestine.

WNT2B deficiency causes a developmental phenotype in human intestine with HIOs showing a decrease in their mesenchymal component and WNT2B-deficient patients showing epithelial disorganization.

**Data Transparency Statement:** All RNA-Seq data will be available through online repository as indicated in Transcript profiling. Any other data will be made available upon request by emailing the study authors.

## Introduction

Canonical wingless-related integration site proteins (Wnts) bind to low density lipoprotein receptor-related proteins (LRP) 5 and LRP6 as well as Frizzled (FZD) receptors on target cells, leading to the dissociation of the AXIN-APC-GSK3β complex and allowing for β-catenin to translocate into the cell nucleus to activate its transcriptional program (reviewed in ^1^). β-catenin signaling is critical for a number of developmental processes including organogenesis and specification of laterality and is also important for stem cell maintenance. Wnt signaling is modulated by the presence of R-spondins, which bind to Leucine-rich repeat-containing G-protein coupled receptor (LGR) 4, LGR5, and/or LGR6 receptors, which impact the cell surface expression of Wnt receptors (reviewed in ^2^). Several Wnt family members play important roles in intestinal homeostasis. WNT3 and WNT2B are canonical Wnts that are expressed in the intestine, which support LGR5^+^ intestinal stem cells (ISCs). In the small intestine, WNT2B is thought to be primarily expressed by PDGFRA^lo^ stromal cells including CD81^+^ trophocytes and FOXL1^+^ telocytes ^3–7^, while WNT3 is primarily expressed in the epithelium by Paneth cells ^8^.

Several lines of evidence suggest that WNT3 and WNT2B are fully redundant in the intestine. First, epithelial WNT3 was not required for small intestinal homeostasis in *Wnt3^Vil-CreERT^*^2^ mice ^9^. Furthermore, WNT2B was the only mesenchyme-expressed Wnt that could support *Wnt3* knock out (KO) organoids in culture ^9^, suggesting these WNT ligands are functionally redundant. In addition, administration of exogenous WNT2B or WNT3 were both able to rescue the intestine in conditional *Wntlss* KO mice, which cannot transport Wnts out of their host cell and are thus functionally Wnt null ^10^. Moreover, mouse intestinal organoids can be supported with either exogenous WNT3 or WNT2B, consistent with their functional redundancy ^9^. Finally, neither mesenchymal- nor epithelial-specific deletion of Porcupine, which is required for palmitoylation of Wnts, led to phenotypic changes in the intestine ^4, 11^.

Despite experimental evidence for redundancy with WNT3 in mice, at least under non-stressed conditions, humans with loss-of-function mutations in *WNT2B* are severely affected by neonatal-onset intractable diarrhea and chronic inflammation of the gastrointestinal tract ^12, 13^. The marked intestinal disease resulting from WNT2B deficiency indicates a critical role in intestinal health and that WNT3 is not fully redundant with WNT2B in humans. To investigate WNT2B’s role during intestinal development and homeostasis, we studied the intestine and colon of *Wnt2b* KO and WT mice as well as patient-derived WNT2B-deficient human intestinal organoids (HIOs) and primary tissues.

## Methods

### Research Ethics

The study was approved by Boston Children’s Hospital Institutional Review Board for the human subjects research (BCH IRB 10-02-0053 and P00027983) in accordance with the Declaration of Helsinki.

### Reagents

A table of reagents used in this study, including antibodies and qPCR primers, is included in Supplementary Table 1 and recipes for media in Supplemental Table 2.

### Mouse model

The ARRIVE guidelines for mouse experimentation were followed. All animal experiments were done in accordance with Boston Children’s Hospital Institutional Animal Care and Use Committee (IACUC, protocols 00001721 and 20-10- 4252). All mice were aged between 2 and 4 months at the time of experiments; male and female mice were used equally. All controls used for experiments were matched littermates. Mice were fed a standard chow diet, bedding was standard animal facility autoclaved pellets, and mice were caged in Optimice cages with Lixit water bottles. For DSS experiments water was placed in free-standing water bottles and the Lixit system was detached. Interventions were performed during the light cycle.

Wnt2b^fl/fl^ mice (C57BL/6N background) were a generous gift from T. Yamaguchi (NCI/NIH). We bred the Wnt2b^fl/fl^ mice with CMV-Cre mice (Jackson labs; B6.C- Tg(CMV-cre)1Cgn/J) to generate Wnt2b^fl/-^ mice, which were then crossed to generate Wnt2b KO mice. Male and female mice were used for experiments at about 3 months of age, and Wnt2b^+/+^ (wild type, WT) or Wnt2b^+/-^ heterozygous (het) matched littermates were used as controls. For all experiments WT and het mice were separately analyzed and no statistically significant differences were found between them, so they are presented as one group (WT) throughout the manuscript.

### Mouse intestinal enteroid and colonoid generation

Mouse organoids were generated as described previously ^14–17^, see Supplemental Methods for full protocol. Enteroids and colonoids were fed with growth media every 2-3 days and passaged every 7 days.

### IFNγ in vitro experiment

Recombinant mouse IFNγ (1,000pg/ml) was added to organoid cultures. The organoids were incubated for 3 days (injury) and the media was replaced without IFNγ and further incubated for another 3 days (recovery).

### Quantitative PCR

Mouse tissue or HIOs were placed into Tri-reagent (ThermoFisher) and RNA was isolated using the Direct-zol kit (Zymo Research) or the RNAEasy mini kit (Qiagen) according to manufacturer’s instructions. RNA was then reversed transcribed into cDNA using a high-capacity cDNA reverse transcription kit and RNAse Inhibitor (ThermoFisher) according to manufacturer’s instructions. Quantitative PCR was performed on a QuantStudio6 Flex (ThermoFisher) using Taqman qPCR Master mix and specific primers. Taqman primers used in analyses are indicated in Supplemental Table 1.

### CD3ε Intestinal Inflammation

Anti-CD3ε (GoInvivo purified, BioLegend) was administered by intraperitoneal injection at a dose of 200μl/25g body weight. The mice were then weighed every 2 days until day 4 post treatment for the injury experiments and again at day 7 for the recovery experiments.

### DSS Colitis Induction

Each experimental cage was supplied with 150ml 2.5% Dextran sodium sulfate (DSS, MPBio) in autoclaved water and changed every two days.

Untreated controls were given autoclaved water. For the injury experiments, mice received DSS for 3 days or 6 days. For the recovery experiments, 2.5% DSS was given until the mice lost 5% of their starting body weight, and then the DSS administration was halted and weight recovery was tracked until the mice had either 20% weight loss or started to regain weight.

### Histological Analysis

For morphologic analysis, intestine or colon was cleaned of feculent material using 4% paraformaldehyde (PFA), and then tissue was sliced longitudinally, rolled and placed in a fixation cassette (Swiss roll method). Tissues were fixed in 4% PFA overnight at 4°C, then transitioned into 70% ethanol. Tissues were embedded in paraffin and sections (4-6μm) were stained with hematoxylin and eosin (H&E) or Periodic Acid Schiff (PAS). Slides were analyzed on an EVOS m7000 (ThermoFisher) at low magnification for imaging intact Swiss rolls (2x) or a Nikon Eclipse E800 microscope with Spot software for higher magnifications.

### Villus Length and Crypt Depth Assessment

Using a Nikon Eclipse E800 microscope with Spot software or a Nikon Eclipse 90i with Nikon Elements software, crypts with a visible lumen were measured from the center base to the level of the crypt-villus transition. Villi were measured from the center of the villus tip to the middle base at the level of the villus-crypt transition in non-oblique sections.

### Immunofluorescence

Slide preparation for immunofluorescence is detailed in Supplementary Methods.

### Generation of WNT2B-deficient iPS line

The human iPS cell lines were established as described in Supplemental Methods. The hiPSCs were cultured in mTeSR1 (StemCell Technologies, 85850) on Matrigel (Corning, 354277)-coated plates and passaged according to the manufacturer’s instructions.

### Human intestinal organoid (HIO) generation

Human iPS cells were differentiated into intestinal organoids as described ^18–20^ with some modifications. Please see Supplemental Methods for full details.

### RNA-Seq

HIOs were placed into Trizol LS (ThermoFisher) followed by isolation of RNA using the Direct-zol kit (Zymo Research) according to the manufacturer’s instructions.

The library was prepared by the Dana Farber Next Generation Sequencing Core Facility using a KAPA library quantification kit (Roche). Sequencing was done on an Illumina NovaSeq 6000 with total RNA.

All samples were processed using a bulk RNA-seq pipeline implemented in the bcbio-nextgen project (bcbio-nextgen,1.2.8-1c563f1). Gene expression was quantified using Salmon (version 0.14.2) ^21^ using the hg38 transcriptome (Ensembl). Differentially expressed genes between the control HIOs and the WNT2B-deficient HIOs were identified using DESeq2 (version 1.30.1) ^22^. The R package clusterProfiler (version 3.18.1) ^23, 24^ was utilized to identify and visualize the enriched Gene Ontology categories. All the analyses performed in R utilized R version 4.0.3.

### Statistical Analyses

For experiments comparing two groups (e.g., control mouse vs KO mouse) Student’s t test was used in PRISM 9 (GraphPad Software). For experiments assessing the impact of treatment conditions (anti-CD3 or DSS) on Control vs KO mice, 2-way analysis of variance (ANOVA) with multiple comparisons and Tukey correction was used in PRISM 9. Data were assumed to be normally distributed, and for outcome variables with highly different standard deviations we used Welch’s correction for t tests. For analyzing weight loss over time, linear regression was used in SAS (SAS Institute). Statistical analysis of RNA-Seq data is described separately in that section. For all experiments p<0.05 was considered significant.

## Results

### Wnt2b KO mice have normal small intestinal histology and differentiation markers

To determine whether *Wnt2b* KO mice have the histologic changes seen in humans with WNT2B deficiency (chronic inflammatory changes with atrophic epithelium)^12, 13^, we examined H&E staining of the duodenum and ileum of *Wnt2b* KO and WT mice. The intestinal architecture was similar between WT and KO mice (Fig 1A). Examination of PAS staining showed normal distribution of Paneth cells and goblet cells in the epithelium (Fig. 1B). To assess for differences in the expression of cell-specific markers in the duodenum, we used qRT-PCR to measure *Lgr5* (ISC), *Lyz1* (Paneth cells), *Muc2* (goblet cells), and *Alpi* (enterocytes) (Fig. 1C). While *Lgr5* expression was significantly decreased in the *Wnt2b* KO mice (about 40% less than controls), the expression of differentiated lineage markers was similar in both mice. As expected, *Wnt2b* was not expressed in *Wnt2b* KO mice (Fig. 1C).

**Figure 1.**
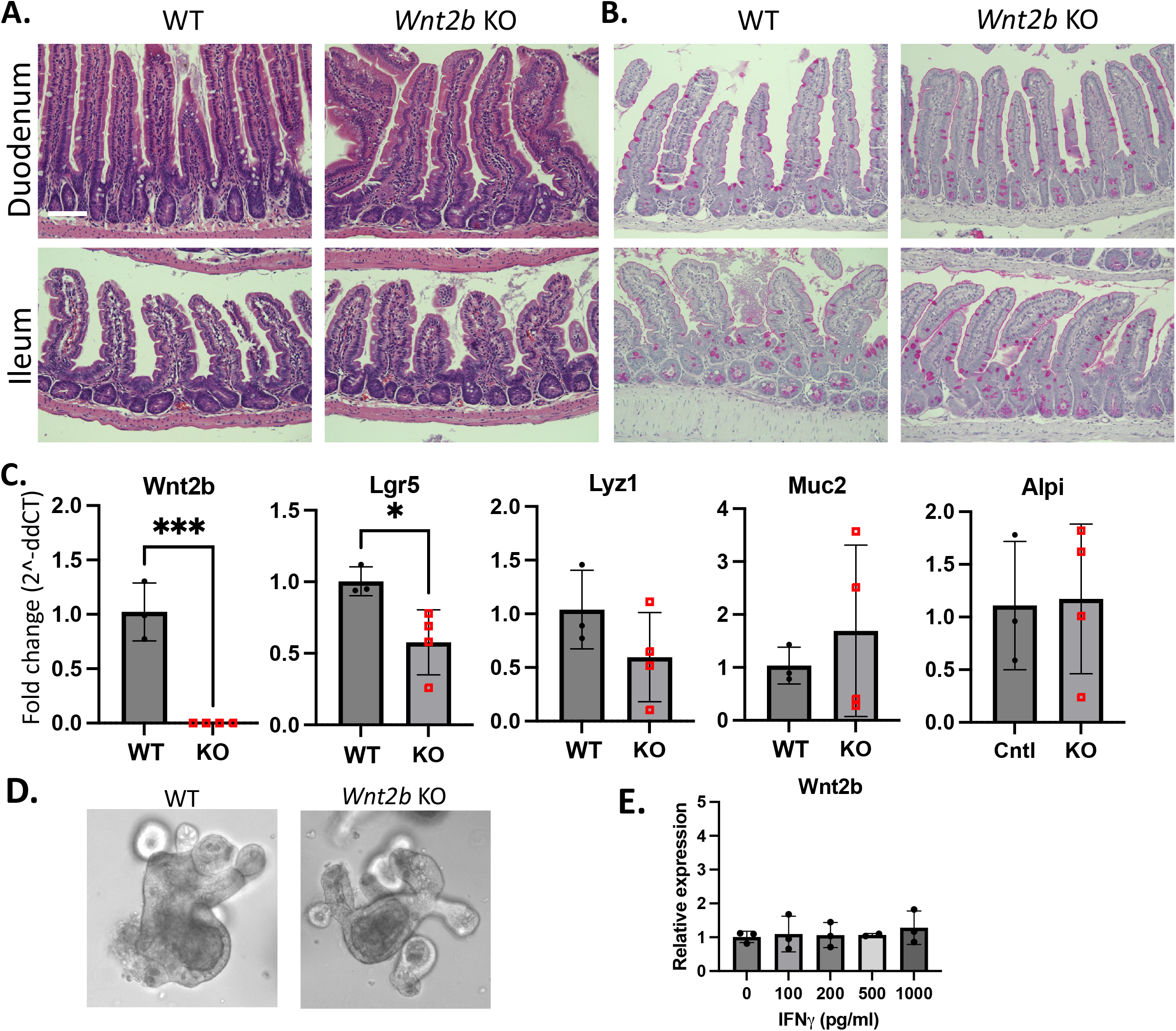
*Wnt2b* KO mice have normal small intestinal histology and lineage marker expression but decreased *Lgr5* expression. **A**. Hematoxylin & eosin staining of small intestine of control versus *Wnt2b* KO mice. **B**. Periodic acid Schiff staining highlighting Paneth cell granules and goblet cells. **C**. Gene expression of *Wnt2b* and epithelial lineage markers by qRT-PCR. **D**. Epithelial small intestinal organoids (enteroids). **E**. Expression of *Wnt2b* by epithelial organoids treated with increasing doses of interferon-γ for 3 days as determined by qRT-PCR. Scale bar denotes 200μm.*p<0.05, ***p<0.001

### Wnt2b KO enteroids are viable and expand normally in standard media

The intestinal mesenchyme is the dominant source of WNT2B production in the intestine, however, others have reported expression of WNT2B from the epithelium in enteroids^25^ and have shown that exogenous WNT2B can support epithelial organoids in culture^9^. Therefore, we wanted to examine whether *Wnt2b* KO organoids would grow *in vitro* in the absence of WNT2B, normally contributed by the mesenchymal milieu *in vivo*. *Wnt2b* KO small intestinal organoids (enteroids) could be established using our standard organoid culture media, which lacks exogenous Wnt ligands (see Supplemental Table 2,) and showed similar morphology to WT controls (Fig. 1D). The organoids expanded similarly to WT enteroids for over 15 passages, indicating normal growth and cell division capacity. We also sought to determine whether *Wnt2b* expression is increased in enteroids under inflammatory conditions by treating the enteroids with interferon-γ (IFNγ), which causes Paneth cell destruction and organoid death ^26^. *Wnt2b* RNA was detected at low levels (qRT-PCR cycle count 35-37) in enteroids but was not impacted by IFNγ dose (Fig. 1E), supporting reports that the epithelium can produce WNT2B, albeit at low levels ^25, 27^. When we treated the enteroids with a dose of IFNγ that caused homogenous organoid injury (1,000pg/mL) for three days, and then allowed the cultures to recover for three days in standard media, there was no difference between WT organoids and *Wnt2b* KO enteroids (Supplemental Fig. 1A) either at the peak injury phase at day 3 or the recovery phase at day 6, with recovering enteroids showing similar numbers of budding crypts. This suggests that WNT2B is not critical for post-injury regeneration.

### Wnt2b KO mice have a normal response to anti-CD3e antibody-induced inflammation

In contrast to humans with WNT2B deficiency, mice with WNT2B deficiency might not show phenotypic changes because they are housed in specific-pathogen free conditions and may lack required inflammatory or infectious drivers of pathology. Similarly, enteroids may not fully recapitulate *in vivo* conditions. Thus, we wanted to challenge the *Wnt2b* KO mouse small intestine to an *in vivo* inflammatory insult using the anti-CD3ε model of small intestinal injury ^28^. In this model, peak injury occurs at day 4 post-injection, so we chose this time point for analysis of tissue morphology. Treating the mice with anti-CD3ε led to similar weight loss in WT and *Wnt2b* KO mice (Fig. 2A). H&E analysis of the small intestine at 4 days post-treatment was also similar (Fig. 2B). Anti- CD3ε antibody treatment did not affect villus length while it did lead to an increase in crypt depth compared to untreated mice, as previously reported ^29^, but there was no difference in crypt elongation between *Wnt2b* KO and WT mice (Fig. 2C, 2D, 2E).

**Figure 2.**
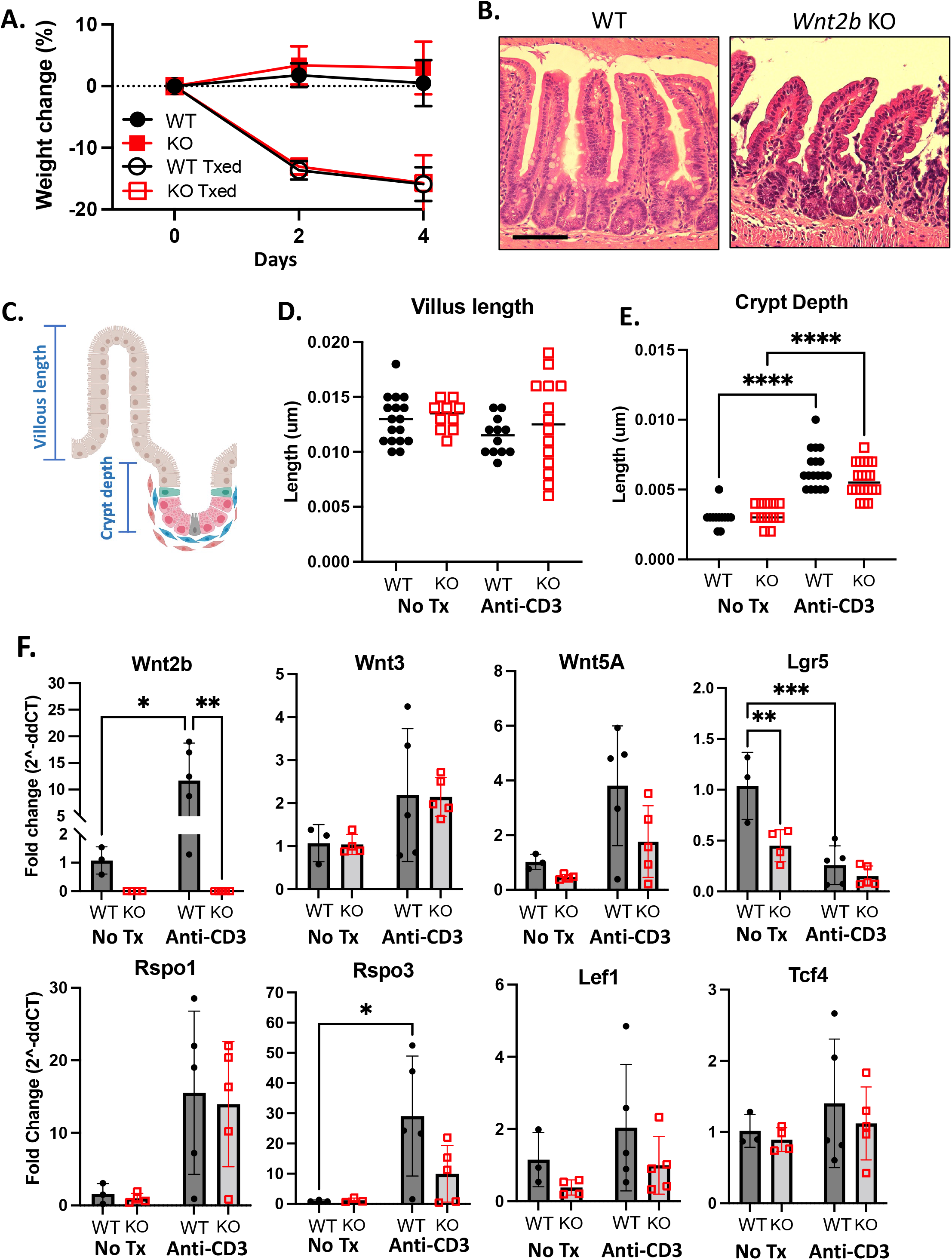
*Wnt2b* KO Mice Have Normal Small Intestinal Response to anti-CD3 Inflammation. **A**. Weight change as a percent of initial weight for control and *Wnt2b* KO mice after injections with anti-CD3ε antibody on day 0 through day 4. **B**. Hematoxylin & eosin staining of small intestines from recovered mice at day 4. **C**. Schematic showing measurements of villus length (in **D**.) and crypt depth (in **E**.) for the treatment groups and controls. **F**. RNA expression of intestinal Wnts, Wnt potentiators, and β-catenin target genes by q-RT-PCR. Scale bar denotes 200μm. *p<0.05, ** p<0.01, ***p<0.001

Because there was no morphologic difference between the KO and WT mice, we next analyzed expression of Wnt ligands, effectors, and downstream targets, to assess whether other intestinal Wnts were increased to compensate for *Wnt2b* deficiency in response to CD3ε treatment (Fig. 2F). We noted a marked increase in *Wnt2b* expression in WT mice in response to anti-CD3ε. Expression of *Rspo3* was also increased in anti-CD3ε treated mice, and *Rspo1*, *Wnt3*, and *Wnt5A* all showed a trend toward higher expression, but no difference in expression between WT and KO mice. In contrast, *Lgr5* expression was significantly decreased in all groups compared to untreated WT mice. Downstream transcriptional targets of β-catenin signaling, *Lef1* and *Tcf4*, were similar regardless of treatment condition or genotype. Analysis of mice treated with a single dose of anti-CD3ε antibody and allowed to recover through 7 days post treatment showed that *Wnt2b* KO mice recovered their weights similarly to WT mice (Supplemental Fig. 2), again suggesting WNT2B is not required for recovery from anti-CD3ε antibody-induced injury in the SI.

### Wnt2b KO colons have normal small intestinal histology and differentiation markers

Next, we examined the colon of *Wnt2b* KO mice by H&E and PAS, and observed no differences between WT and *Wnt2b* KO mice (Fig. 3A and 3B). qRT-PCR for colonic epithelial markers revealed a significant decrease in expression of *Lgr5*, roughly an order of magnitude lower than in control mice, while markers of epithelial cells, including *Alpi* (enterocytes) and *Muc2* (goblet cells), were similar to WT mice (Fig 3C). In addition, colon organoids (colonoids) generated from WT and *Wnt2b* KO mice grew similarly (Fig. 3D) in standard colon media (see Supplemental Table 2) and expanded normally for >15 passages. WT colonoids had stable expression of *Wnt2b* when treated with increasing doses of IFNγ (Fig. 3E). Moreover, challenging the colonoids with IFNγ led to a similar injury between WT and *Wnt2b* KO colonoids at day 3 and robust recovery in both groups at day 6 (Supplemental Fig. 1B), suggesting that WNT2B is not required for recovery from IFNγ-induced injury in colonoids.

**Figure 3.**
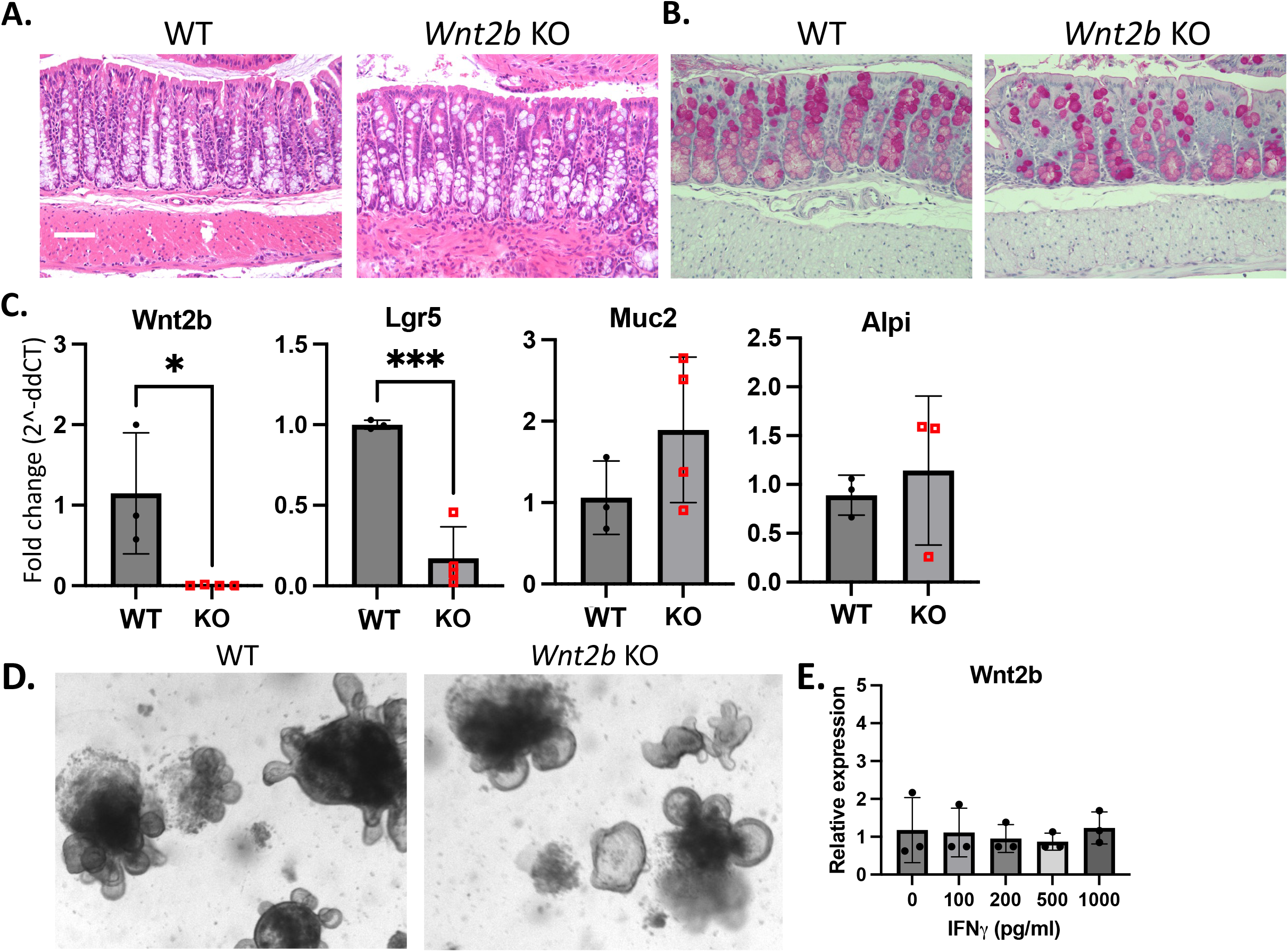
*Wnt2b* KO mice have normal colon histology and lineage marker expression but decreased *Lgr5* expression. **A**. Hematoxylin & eosin staining of colon of control versus *Wnt2b* KO mice. **B**. Periodic acid Schiff staining highlighting Paneth cell granules and goblet cells. **C**. Gene expression of *Wnt2b* and epithelial lineage markers by qRT-PCR. **D**. Epithelial colon organoids (colonoids). **E**. RNA expression of *Wnt2b* by control colonoids with increasing doses of interferon-γ for 3 days. Scale bar denotes 200μm. *p<0.05, ***p<0.001

### Wnt2b KO colon has increased sensitivity to dextran sodium sulfate colitis (DSS)

Next, we examined the response of the WNT2B-deficient mouse colon to inflammation *in vivo*. Analysis of *Wnt2b* KO mice treated with DSS showed progressive weight loss by day 5 and 6 of treatment compared with WT mice (Fig 4A). In fact, the experiment had to be stopped at day 6 because the *Wnt2b* KO mice had reached the prespecified early experiment termination endpoint of twenty percent weight loss by day 6. Analysis of colon size was similar between the *Wnt2b* KO and WT mice at day 6 (Fig. 4B); however, histological analysis of PAS staining showed loss of goblet cells in the DSS-treated *Wnt2b* KO mice, while H&E staining demonstrated crypt loss and lymphocytic infiltration (Fig. 4C). Blinded histologic scoring using published metrics for DSS colitis ^30^ confirmed a significantly worse injury in *Wnt2b* KO mice at day 6 with increased goblet cell loss, increased crypt loss, and increased submucosal inflammatory infiltrates but no change in muscle thickness or evidence for crypt hyperplasia (Fig. 4D, see Supplemental Table 3 for scoring parameters). These data show that WNT2B is important for resistance to colitis in the mouse.

**Figure 4.**
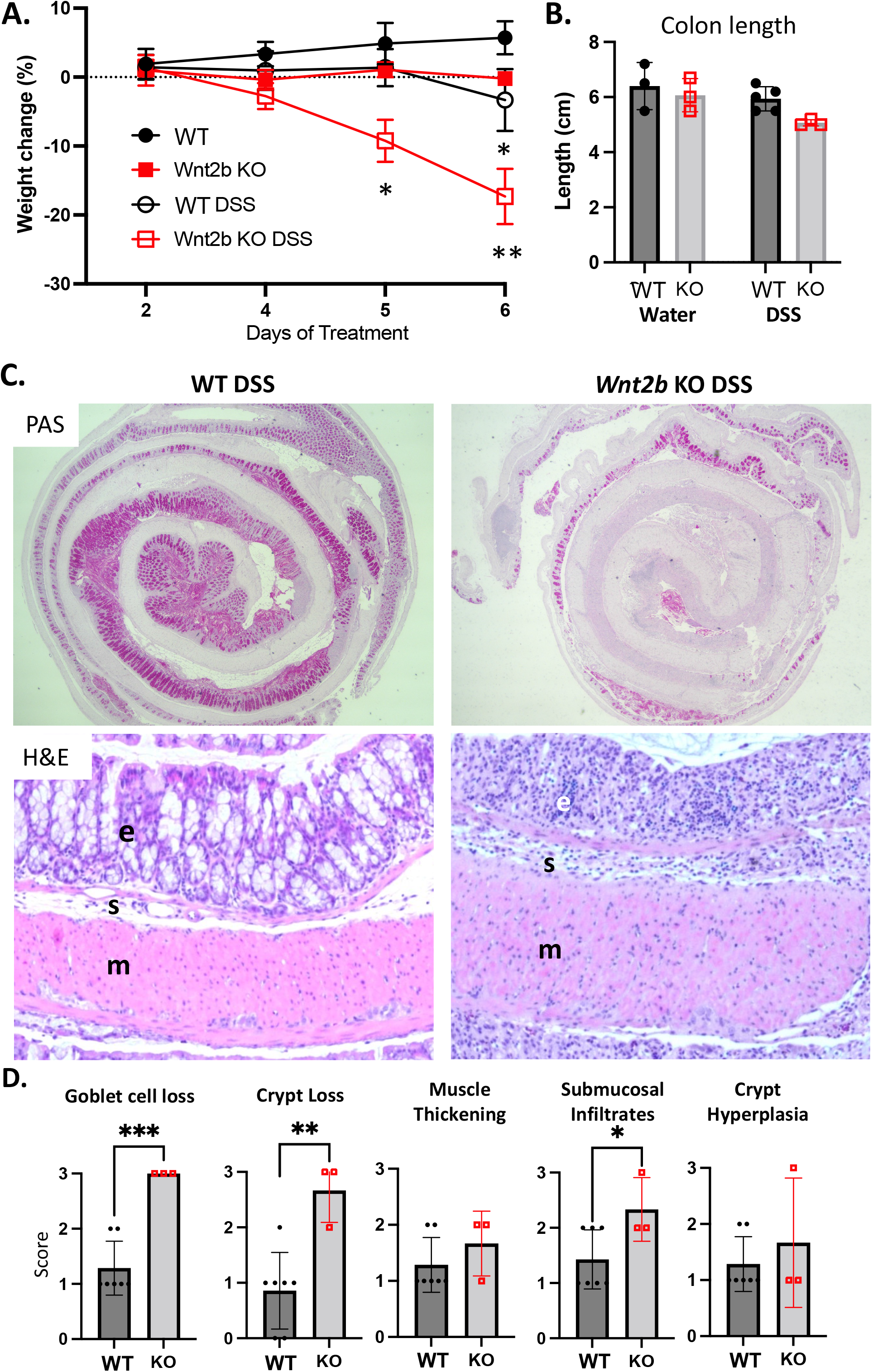
*Wnt2b* KO Mice Have Increased Susceptibility to DSS Colitis. **A**. Weight change as a percent of initial weight for control and *Wnt2b* KO mice treated with 2.5% DSS in drinking water for 6 days. **B**. Colon lengths on day 6. **C**. PAS and H&E staining of colon after 6d DSS. e - epithelium, s - submucosa, m - mesenchyme. **D**. Blinded histologic assessment of DSS colitis features for control vs KO mice after 6d DSS. *p=<0.05, **p=<0.01, ***p=<0.001

### Wnt2b KO mice demonstrated similar intestinal proliferation compared to WT mice

Assessment of *Wnt2b* expression in the colon at day 6 of DSS treatment showed no differences between treated and untreated WT mice (Fig. 5A). In contrast, markers of intestinal stem cells, *Lgr5* and *Ascl2*, were significantly decreased in WT mice treated with DSS by day 6 compared to untreated mice, which were similar to expression levels in *Wnt2b* KO mice at baseline. Expression of *Atoh1*, which has been shown to be important for repopulating the ISC niche after DSS colitis through dedifferentiation of secretory precursor cells ^31^, was similar between *Wnt2b* KO and WT mice, indicating that this pathway remains intact in the absence of WNT2B. Analysis of proliferation, as measured by Ki67 staining, was similar in untreated WT and *Wnt2b* KO mice and decreased after 6d of DSS treatment in both groups (Fig. 5B).

**Figure 5.**
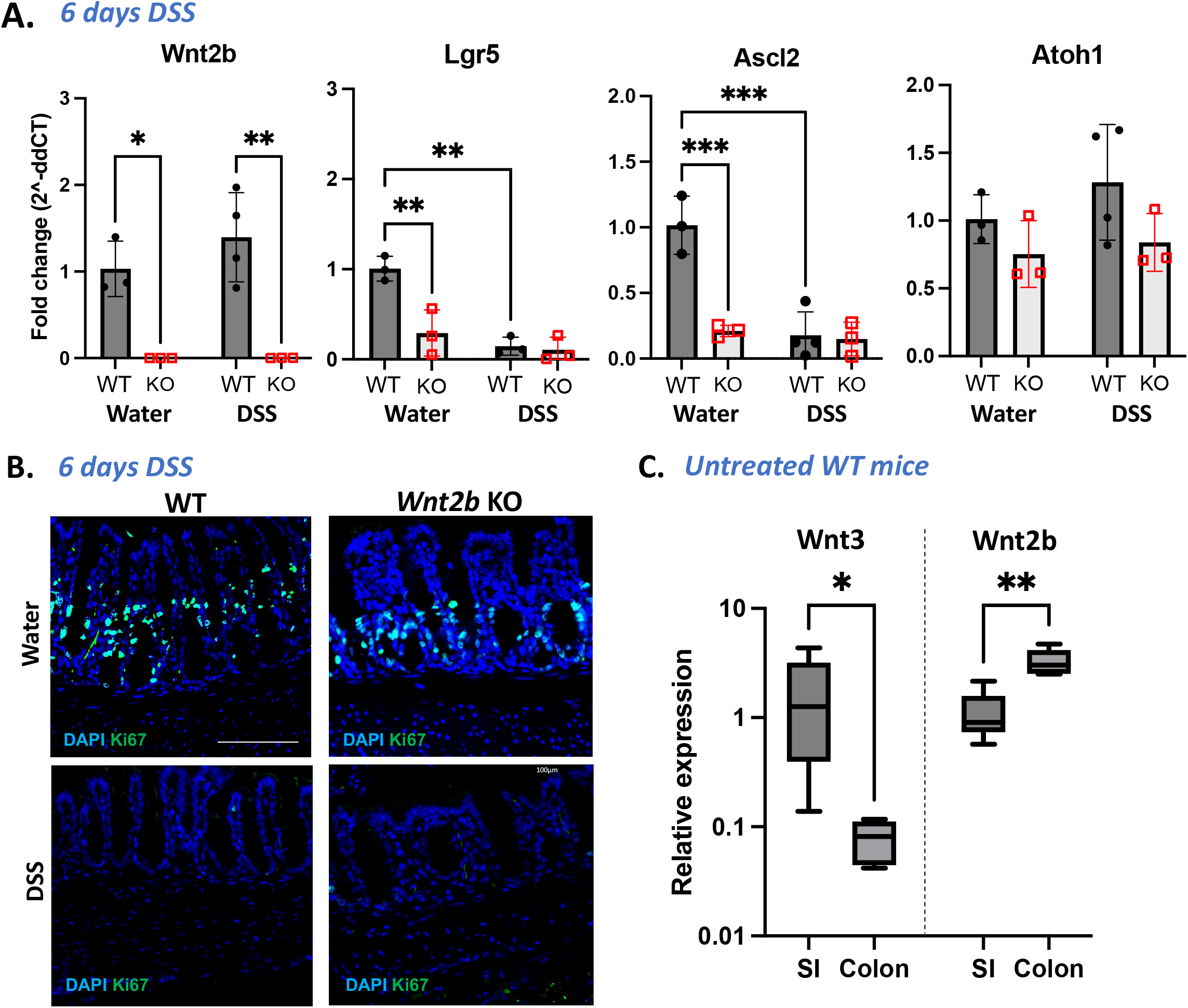
*Wnt2b* KO mice show faster time to DSS-induced injury but preserved proliferation. **A**. Expression of *Wnt2b*, *Lgr5*, *Ascl2*, and *Atoh1* in DSS treated and control mice after 6 days of 2.5% DSS or water by qRT-PCR. **B**. Expression of Ki67 (green) and DAPI (blue) in DSS and water treated mice after 6 days. **C**. Expression of *Wnt2b* and *Wnt3* in whole thickness intestinal tissue in small intestine (SI) and colon from untreated wild type mice. Scale bar = 100μm. *p<0.05, ** p<0.01, ***p<0.001

To assess whether *Wnt2b* KO mice have increased sensitivity to colitis or impaired recovery from colitis compared to controls, we treated mice with DSS until the first day that each individual mouse lost at least 5% of their starting body weight (Supplemental Fig. 3, indicated by cyan shading in the figure). WT mice took more time to lose at least 5% body weight, but the trajectory of the two groups was similar after this initial lag in the control mice: the slopes of weight loss in both groups were not different (p=0.46). Several mice in each group had to be euthanized for excessive weight loss despite stopping the DSS several days prior, while several mice in each group regained weight. This pattern suggests that the increased susceptibility of *Wnt2b* KO mice to DSS colitis is based on the initial resistance to injury, rather than the rate of development of colitis or the ability to recover from injury.

It has been shown that epithelial proliferation and *Lgr5* expression initially increases around day 3 of DSS treatment in WT mice, before falling below baseline levels by day 7 ^32^, so we decided to evaluate proliferation after 3 days of DSS treatment. Ki67 staining was similar between *Wnt2b* KO and WT mice at 3 days post DSS, (Supplemental Fig 4A). We also evaluated differences in stem cell marker expression between *Wnt2b* KO and WT mice after 3 days of DSS (Supplemental Fig. 4B). DSS-treated WT mice trended toward higher expression of stem cell markers than untreated WT mice at Day 3 of DSS (Lgr5 p=0.07), while *Lgr5* and *Ascl2* levels remained unchanged in *Wnt2b* KO mice with and without DSS. Wnt2b KO mice treated with DSS had significantly decreased levels of *Lgr5* at day 3 compared to WT mice. Others have shown that *Bmp4* and *Smad4* expression drive this early proliferative response to DSS, and *Wnt2b* KO mice increased expression of these genes in the setting of colitis (Supplemental Fig. 4B; while not significant by ANOVA with the WT groups included (p=0.13), the difference between *Wnt2b* KO untreated and *Wnt2b* KO DSS groups was significant if analyzed by direct comparison using Mann-Whitney U test (p=0.05)). This suggests that Wnt2b KO mice are able to upregulate *Bmp4* and *Smad4* expression but cannot increase *Lgr5* expression in the early injury phase. This could be either due to the lower *Lgr5* expression at baseline in the absence of WNT2B, or because WNT2B is required for the downstream response to BMP4 and SMAD4.

### Wnt2b is expressed at relatively higher levels in the colon than the SI, in contrast to Wnt3

Previous studies have shown that WNT3 is able to support ISCs *in vitro*, and that WNT2B and WNT3 are functionally redundant ^9–11^. To begin to understand why the mouse colon was more sensitive to loss of WNT2B than the small intestine, we examined the relative expression of *Wnt3* and *Wnt2b* in the small intestine and colon of WT mice. *Wnt3* expression was significantly lower in the colon compared with the SI (Fig. 5C). Conversely, *Wnt2b* expression was significantly higher in the colon than the SI. These data raise the possibility that the colon has a more significant phenotype in the absence of WNT2B, because it produces less WNT3 and thus may be more dependent on the local production of WNT2B.

### Human Wnt2b-deficient iPS-derived human intestinal organoids show abnormal epithelial development

Humans with WNT2B deficiency have a profound presentation and are affected from the neonatal period in both the small intestine and colon ^12, 13^. We therefore sought to examine the developmental impact of WNT2B deficiency in human tissues using patient-derived iPS to generate human intestinal organoids (HIOs), which contain both epithelial and mesenchymal layers and express immature, fetal-like markers of intestinal development ^20^ (Fig. 6A). Using previously published methods to induce differentiation into definitive endoderm and then into intestinal spheroids ^19, 20^, we generated HIOs from WNT2B-deficient patient iPS and from control iPS lines. WNT2B- deficient HIOs were morphologically distinct from controls and had a smaller size and a shifted ratio of central spheroid tissue to external tissue (Fig. 6B, 6C). Confocal microscopy of whole-mount immunofluorescence staining for mesenchymal marker vimentin and epithelial marker E-cadherin consistently showed abnormal organization of the epithelium in the WNT2B-deficient HIOs compared with controls. The E-cadherin staining was particularly notable for cystic structures in the epithelium and DAPI nuclear staining in the E-cadherin expressing epithelial regions indicated abnormal organization of epithelial nuclei, which were not in the typical linear, closely approximated rows (Fig. 6D; Supplemental Fig. 5).

**Figure 6.**
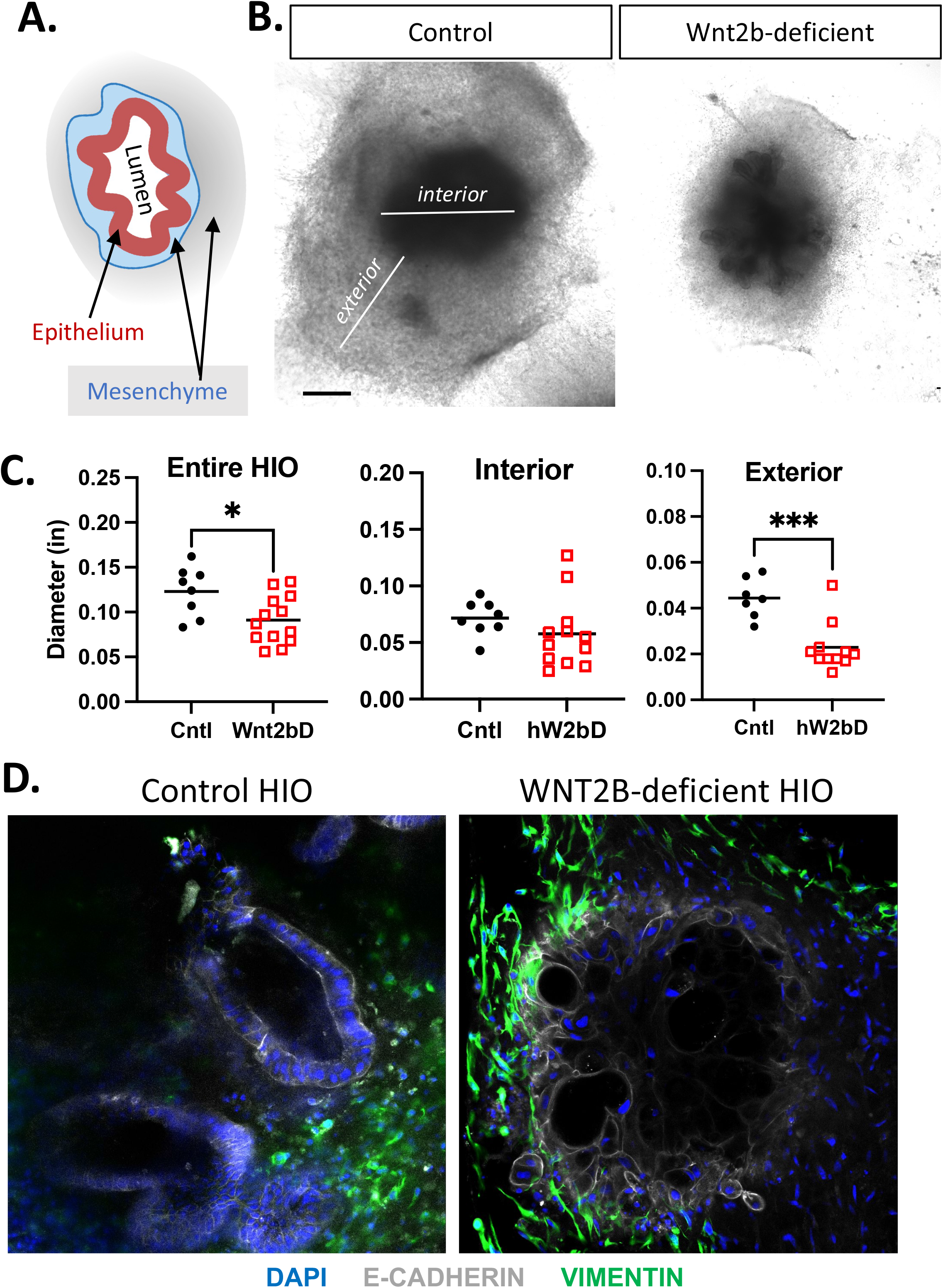
WNT2B-deficient human intestinal organoids have abnormal epithelial organization. **A**. Schematic of an HIO showing approximate regions of epithelium, mesenchyme, central lumen. **B**. Brightfield microscopy of control and WNT2B-deficient HIOs at 28 days in culture. **C**. Measurements of the total diameter, interior and exterior of control versus WNT2B-deficient HIOs. **D**. Immunostaining of HIOs epithelial (E- cadherin), mesenchyme (vimentin) and DNA (DAPI). Scale bar = 50μm. *p<0.05, ***p<0.001

Next, we performed RNA-Seq analysis to assess for differential gene expression between the WNT2B-deficient and control HIOs. Principal component analysis showed that variance was driven by sample type (Supplemental Fig. 6A) and hierarchical clustering also demonstrated that data clustered by experimental replicates (Supplemental Fig. 6B). WNT2B-deficient HIOs showed significant differences in gene expression in muscle-related and extracellular matrix genes as well as in genes related to cell junction and cell adhesion molecules (Fig. 7A). We next assembled heatmaps of curated gene sets specific for the human intestinal mesenchyme (GO:0060485) and epithelium (GO:0060576) (Fig 7B), which showed that mesenchymal genes were upregulated in the WNT2B-deficient HIOs. The epithelial genes, in contrast, did not show a consistent trend of gene expression patterns in WNT2B-deficient HIOs compared to controls.

**Figure 7.**
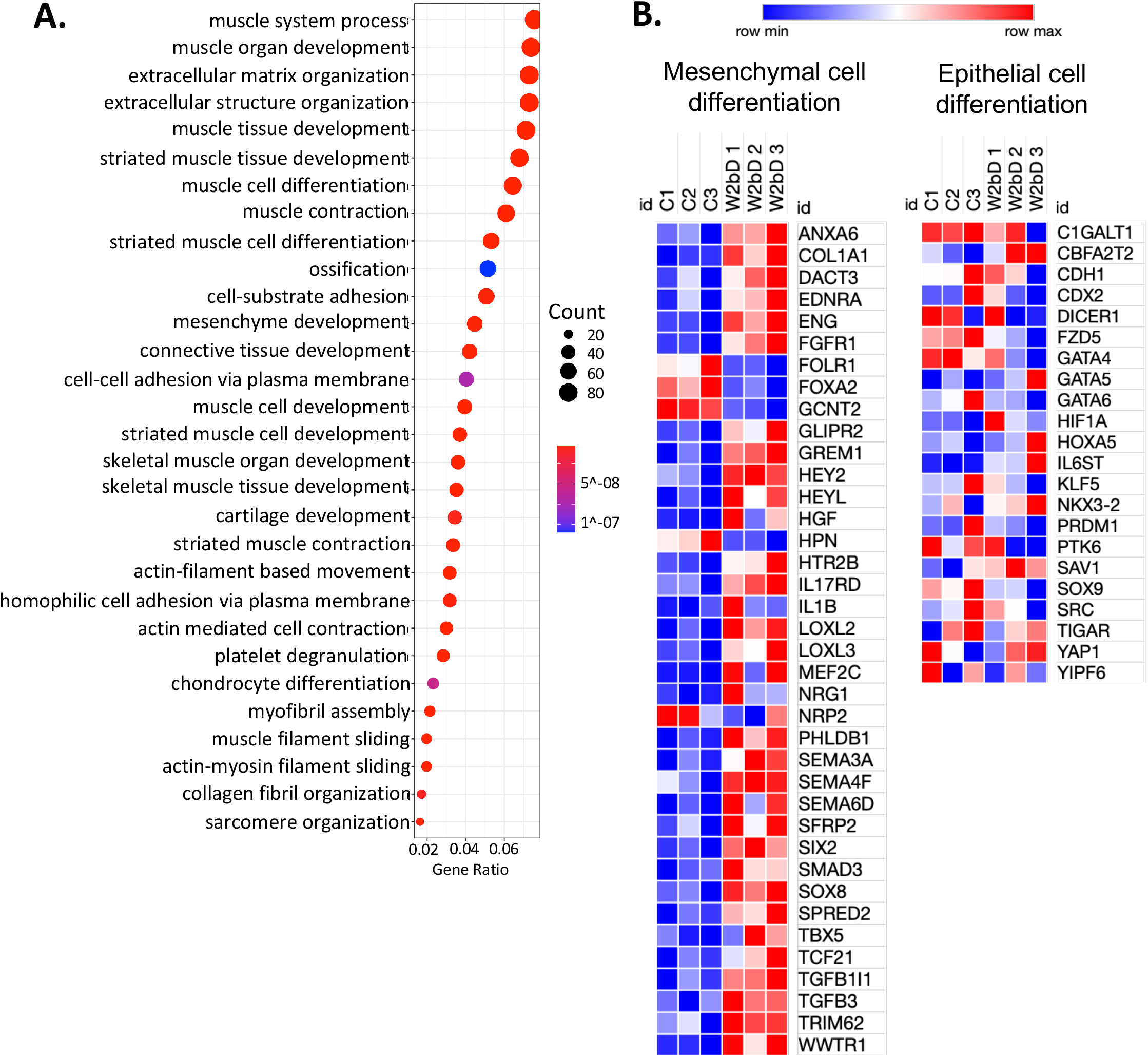
WNT2B-deficient human intestinal organoids have increased expression of mesenchymal genes. **A**. Dot plot of the most differentially expressed gene sets in WNT2B-deficient versus control HIOs. **B**. Heat map of gene sets for mesenchymal and epithelial genes in control (C1/2/3) and WNT2B-deficient (W2bD1/2/3) HIOs.

### Intestinal tissue from patients with WNT2B deficiency is disorganized and shows decreased proliferation

Because the HIOs demonstrated abnormal developmental organization of the epithelium, we re-examined tissue specimens from WNT2B-deficient patients. E-cadherin staining of WNT2B-deficient small intestine (Supplemental Fig. 7A) and colon (Supplemental Fig. 7B) showed normal epithelial organization at the cellular level but highlighted an abnormal epithelial structure at the tissue level along with diminished crypt numbers. Interestingly, biopsies from patients demonstrated diminished staining for Ki67 in both the colon and the small intestine, in contrast to our findings in the *Wnt2b* KO mice. This indicates that WNT2B-deficient patients also have abnormal epithelial organization, similar to the HIOs, along with diminished epithelial proliferation.

## Discussion

We previously reported on human patients with WNT2B deficiency, who have severe intestinal disease form an early age ^12, 13^. This was in contrast to prior studies, all performed in mouse models or epithelial organoid systems, that concluded that WNT2B was not essential and was fully redundant with WNT3 ^9–11, 33^. The present study finds that this discrepancy is partly explained by the fact that loss of WNT2B causes a more profound phenotype in humans than in mice. Furthermore, mice do have a phenotype when WNT2B is deficient, but it requires an injury stimulus, and even then, it is limited to the colon. It will be important to consider the role of injury stimuli in mouse studies designed to evaluate Wnts and other ISC supportive factors moving forward.

The finding that *Lgr5* expression was diminished both in *Wnt2b* KO mouse SI and colon at baseline (Fig. 1C and 3C), while RNA expression of differentiated lineage markers and histology was not different, suggests that ISC are decreased in these mice but not diminished below the threshold required to maintain homeostasis under non-stressed conditions. Inflammatory injury with anti-CD3ε antibody, which targets the SI, did not result in any differences between *Wnt2b* KO and WT mice despite causing a reduction in ISC markers in both mouse types (Fig. 2). In contrast, DSS treatment, which targets the colon, led to robust weight loss and earlier death along with increased histologic abnormalities in the *Wnt2b* KO versus WT mice (Fig. 4). While WT mice showed increased expression of ISC marker genes at day 3 in the early response to DSS, *Wnt2b* KO mice started with lower expression of ISC markers an expression did not increase during the early response to injury. BMP4 and SMAD4 were previously shown to be essential for this initial increase in ISC gene expression ^32^, and *Wnt2b* KO mice were able to increase expression of these genes, indicating that the inability of *Wnt2b* KO mice to respond to early injury was not due to a failure to engage this pathway. (Direct comparison of the *Wnt2b* KO untreated vs *Wnt2b* KO DSS treated groups gave a p value of 0.05 by Mann-Whitney U test, in contrast to the ANOVA multiple groups comparison data in the graph.) *Wnt2b* KO mice also had similar rates of proliferation to WT mice, as measured by Ki67 staining, supporting the ability of *Wnt2b* KO mice to respond with epithelial proliferation after injury. In summary, WNT2B deficiency results in an enhanced initial injury, potentially due to the lower numbers of ISC at baseline, or inability of these ISC to respond to increases in BMP/SMAD signaling.

The increased susceptibility of *Wnt2b* KO mice to DSS colitis compared to anti- CD3ε enteritis may be due to the difference in the relative expression of *Lgr5*, and presumably different numbers of ISC at baseline. *Wnt2b* KO mice had about a 50% decrease in *Lgr5* expression in the small intestine compared with WT mice compared to a 90% reduction in the colon in *Wnt2b* KO mice compared to WT (Fig. 1C and 3C).

Further, the difference in ISC marker expression in the absence of WNT2B in the small intestine may be linked to higher relative expression of *Wnt3* in the small intestine than in the colon (Fig. 5C) as supported by prior studies showing that both WNT2B and WNT3 can support ISCs ^9–11^. Increased *Wnt3* in the small intestine is not altogether surprising since the colon does not contain Paneth cells, which are the primary source of *Wnt3* in the intestine. It was more surprising that relative *Wnt2b* expression in colon was significantly higher than in the small intestine. To our knowledge this has not previously been examined and may have important implications for diseases which differentially impact the colon over the small intestine.

The prevailing dogma has been that *Wnt2b* is expressed solely by the mesenchyme, however, recent studies have shown that epithelial only organoids can produce *Wnt2b* following exposure to *Escherichia coli* secreted protein (Esp) or to radiation ^25, 27^. Similarly, we saw expression of *Wnt2b* in epithelial only enteroids, however we did not see an increase following IFNγ-induced injury. It remains unknown which cell types are responsible for epithelial production of *Wnt2b*.

Remarkably, WNT2B deficiency did not impact recovery in response to the various injury models (IFNγ *in vitro* or anti-CD3ε or DSS *in vivo*) in mice (Supplemental Figs. 1, 2, and 3), possibly due to either intact dedifferentiation pathways or activation of reserve stem cell populations. This suggests that *Wnt2b* is not particularly critical for replenishing the ISC population after injury, at least in mice. In support of this, *Atoh1* expression, which has been demonstrated to be important for replenishing the Lgr5^+^ ISC population during DSS colitis ^31^, was not significantly different between WT and *Wnt2b* KO mice, even during the peak of DSS injury.

WNT2B-deficient patient-derived HIOs showed abnormally organized epithelial structures and demonstrated enhanced expression of mesenchymal transcripts compared to controls (Figs. 6 and 7). WNT2B has been shown to be important in regulating intestinal cell fate determination during the process of epithelial to mesenchymal transition (EMT) ^34^. EMT occurs when a normally polarized epithelial cell begins to take on characteristics of mesenchymal cells, leading to degradation of the basement membrane and subsequent migration away from the typical epithelial position. While EMT is perhaps best described in cancer models, it is also a fundamentally important processes in normal organogenesis ^35^. Target genes associated with EMT including Snail (SNAI1) and Slug (SNAI2) ^36^ were not different in WNT2B-deficient HIOs versus controls. Collagen IV and laminin genes involved in maintaining basement membrane, which is altered in EMT, were also similar between WNT2B-deficient and control HIOs.

The differential response to WNT2B deficiency in the mouse colon versus the small intestine, as well as the relatively increased expression of *Wnt2b* in the colon, hint at the possibility that these WNT2B-producing mesenchymal populations may also differ between the colon and small intestine. A recent publication investigating this question noted a symmetric distribution of *Wnt2b* in the mouse small intestine and the colon, with *Wnt2b* extending the full length of the colonic crypt and the crypt and villus in the SI ^7^.

The total amount of *Wnt2b* was not quantified in these assays, however. The study did not specifically address whether the WNT2B-producing cell populations varied by intestinal region. There may be important clinical implications of increased relative stromal expression of WNT2B in the colon versus the SI, particularly for colon-specific disorders including ulcerative colitis and colon cancer.

Wnt2b KO mice demonstrated similar expression of Ki67 compared with controls (Fig. 5B, Supplemental Fig 4A), indicating that proliferative ability is intact. Functional data in the mice showing that they can recover from either small or large intestinal injury similarly to wild type mice supports intact proliferative responses. In contrast, human small and large intestinal biopsy samples showed diminished Ki67 compared with control subject biopsies from these tissues (Supplemental Fig 7). One potential explanation for this seeming discrepancy is that the patients are more impacted by chronic inflammation.

It is well known that mice and humans have important biological differences, but the exact nature and context of these differences is not always well described. The fact that WNT2B plays an important developmental role in humans, but not in mice, has important implications for intestinal developmental biology. Recapitulating early human intestinal development, for the purpose of regenerative medicine, as an example, will require adequate sources of WNT2B, while this component is not critical for mouse systems. Conclusions made about the relative roles of human intestinal Wnts based on mouse model experiments will need to re-consider this new finding.

## Limitations

This study was limited by the lack of well-validated antibodies to many of the key proteins we investigated, including WNT2B, WNT3, and LGR5. Because of this, we relied heavily on RNA expression data. Certainly, RNA expression is not always reflective of fluctuations at the protein level, and this is an important caveat to our conclusions. This may also impact our conclusions regarding decreased LGR5 expression and presumably diminished stem cell numbers, however the pattern of Ki67 staining combined with the RNA expression data suggest a stem cell mechanism may underlie the defects observed in WNT2B-deficient patients. We also relied on RNA expression when evaluating lineage differentiation, however goblet cells, enterocytes, and Paneth cells are directly visible on PAS staining and the histology supports this conclusion.

Another limitation of the study was slight variability in generating HIOs between batches. To control for inter-batch variability, we ran two separate batches of RNA-Seq and confirmed that the most differentially expressed genes referenced in the paper were common between both sets.

## Conclusion

In sum, WNT2B plays a critical role in early epithelial development in humans while *Wnt2b* KO mice show normal epithelial architecture under normal conditions. Loss of WNT2B in humans and mice leads to diminished *Lgr5* expression, suggesting a decrease in ISC number. Loss of WNT2B in mice caused increased susceptibility to DSS colitis, which may be explained by a relatively higher baseline expression of *Wnt2b* in the colon than the small intestine in mice. Further work should aim to understand the differences between the small intestine and the colon regarding WNT2B, and to further clarify the mechanisms by which WNT2B directs epithelial and stem cell development.

## Abbreviations

DSS: dextran sodium sulfate
GI: gastrointestinal
HIO: human intestinal organoid
iPS: inducible pluripotent stem cells
KO: knock out
Lgr: leucine-rich repeat-containing G-protein coupled receptor
Lrp: low density lipoprotein receptor-related protein
Wnt: wingless-related integration site protein
WT: wild type
SI: small intestine

## Author Contributions

All authors read and critically reviewed the manuscript and approved of the final manuscript. In addition:

AEO conceptualized the study, conceptualized and performed or supervised all of the experiments, and wrote the manuscript.

SR assisted with DSS colitis experiments including qPCRs, fluorescence microscopy, and flow cytometry.

CA performed organoid-based experiments and qPCR analyses.

WQ managed iPS and human intestinal organoid culture.

CE managed the mouse colony and performed qPCR experiments and analyses.

RK performed RNA-Seq bioinformatics analysis and statistics.

JL initially established the Wnt2b KO mouse line.

AS and NR generated iPS from patient fibroblasts and technical assistance with HIO culture.

JDG obtained and processed human intestinal biopsy specimens.

JaRT provided patient specimens and clinical expertise.

DLC provided important intellectual contributions to the project conceptualization and experimental design.

JeRT made important intellectual contributions to the experimental design and interpretation.

PBA provided expertise on genetics and orphan disease and support for study initiation.

MH provided resources and technical expertise on HIO/HCO culture.

DTB provided topical expertise on intestinal biology and specific support for organoid cultures.

## Acknowledgements

The authors are grateful to the patients and families who consented to submit research specimens that were used in this work. We also thank Henry Feldman, Ph.D. for his statistical expertise and advice.

