## Supplemental Figures for "WNT2B Deficiency Causes Increased Susceptibility to Colitis in Mice and Impairs Intestinal Epithelial Development in Humans"

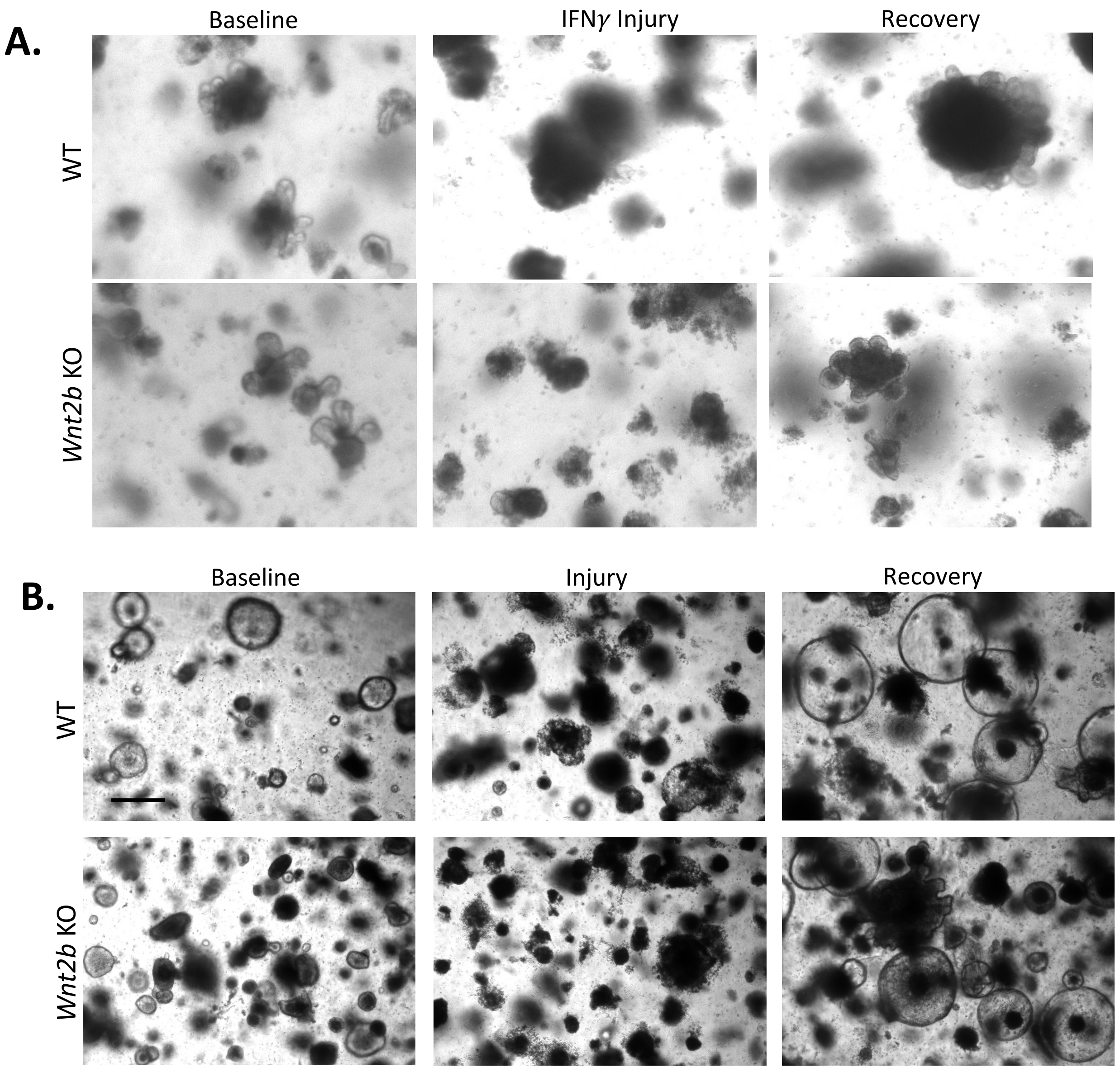


**Supplemental Figure 1. Murine Wnt2b KO enteroids and colonoids can recover from injury**. A. WT and *Wnt2b* KO small intestinal enteroids at baseline (left panel), treated with interferon-γ for 3 days (injury, middle panel), and then allowed to recover for 3 days after IFNγ stopped (recovery, right panel). B. WT and *Wnt2b* KO colonoids at 4x magnification at baseline (left panel), treated with IFNγ for 3 days (injury, middle panel), and then allowed to recover for 3 days after IFNγ stopped (recovery, right panel). Images at 4X on an EVOS XL.

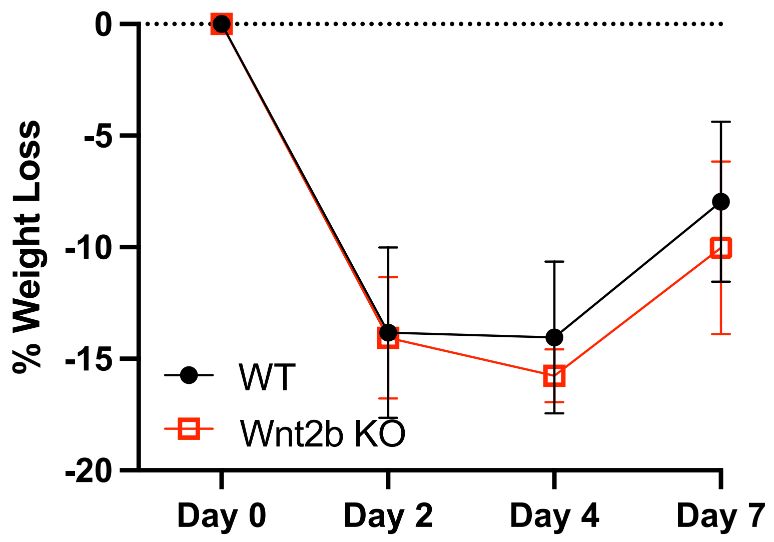


**Supplemental Figure 2. Anti-CD3*ε* mouse experiment 7 days post injection.** WT and *Wnt2b* KO mice were injected with anti-CD3*ε* and weights were followed through 7 days post-injection. Symbols represent the mean weight loss for animals in that group as a percentage of starting weight and error bard indicate standard deviation.

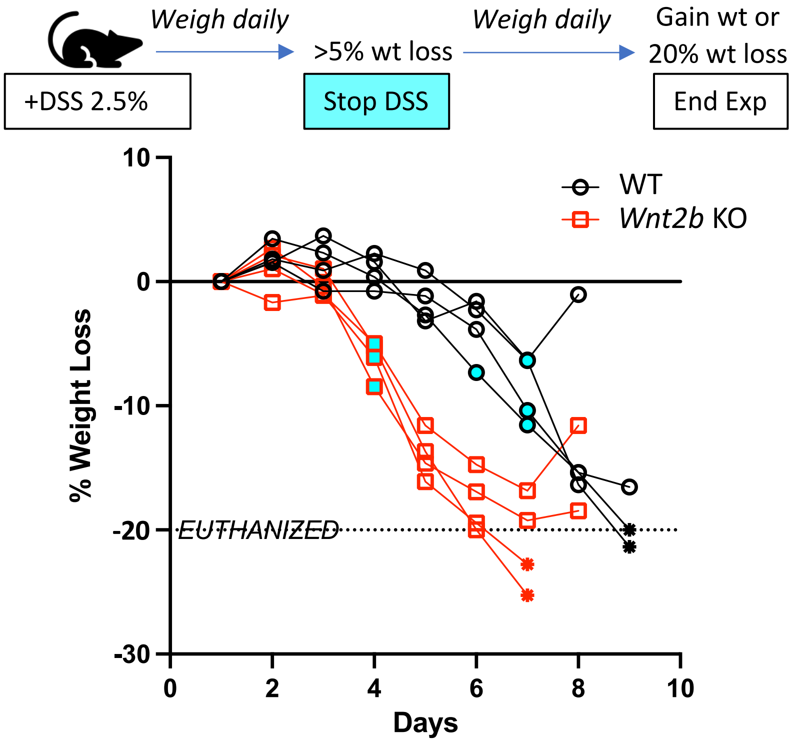


**Supplemental Figure 3.** Weight loss as a percentage of starting weight in mice treated with DSS only until they lost 5% or more of initial weight (cyan dot). Points after this were not treated with DSS. An asterisk here indicates a mouse that was euthanized for excessive weight loss (>20% of starting weight, see dotted line).

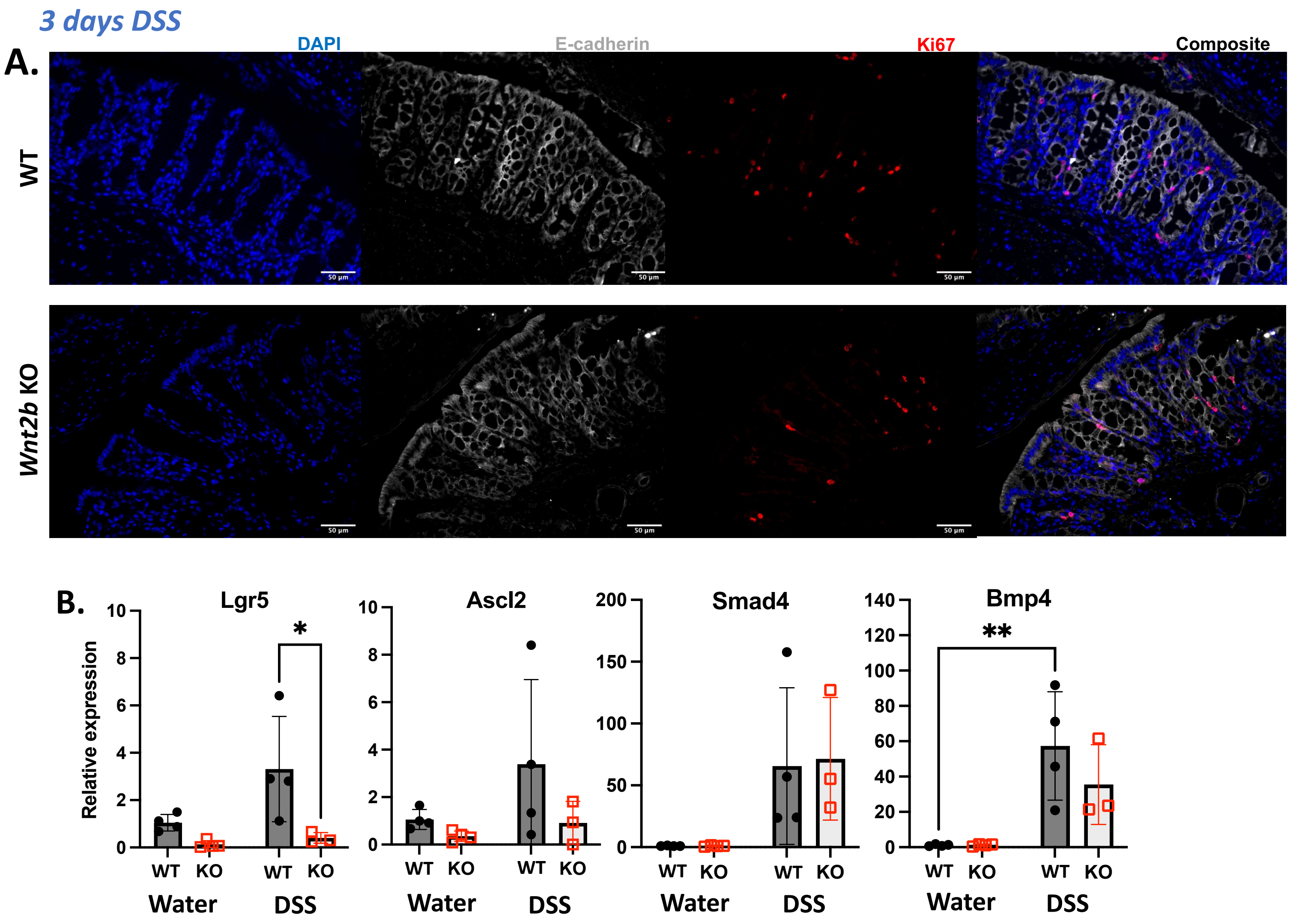


**Supplemental Figure 4. Proliferation and ISC marker expression after 3d DSS.** A. Immunofluorescence staining for Ki67 shows even distribution of Ki67 staining between WT and KO mice. B. Expression of stem cell markers by qRT-PCR. *p=<0.05.

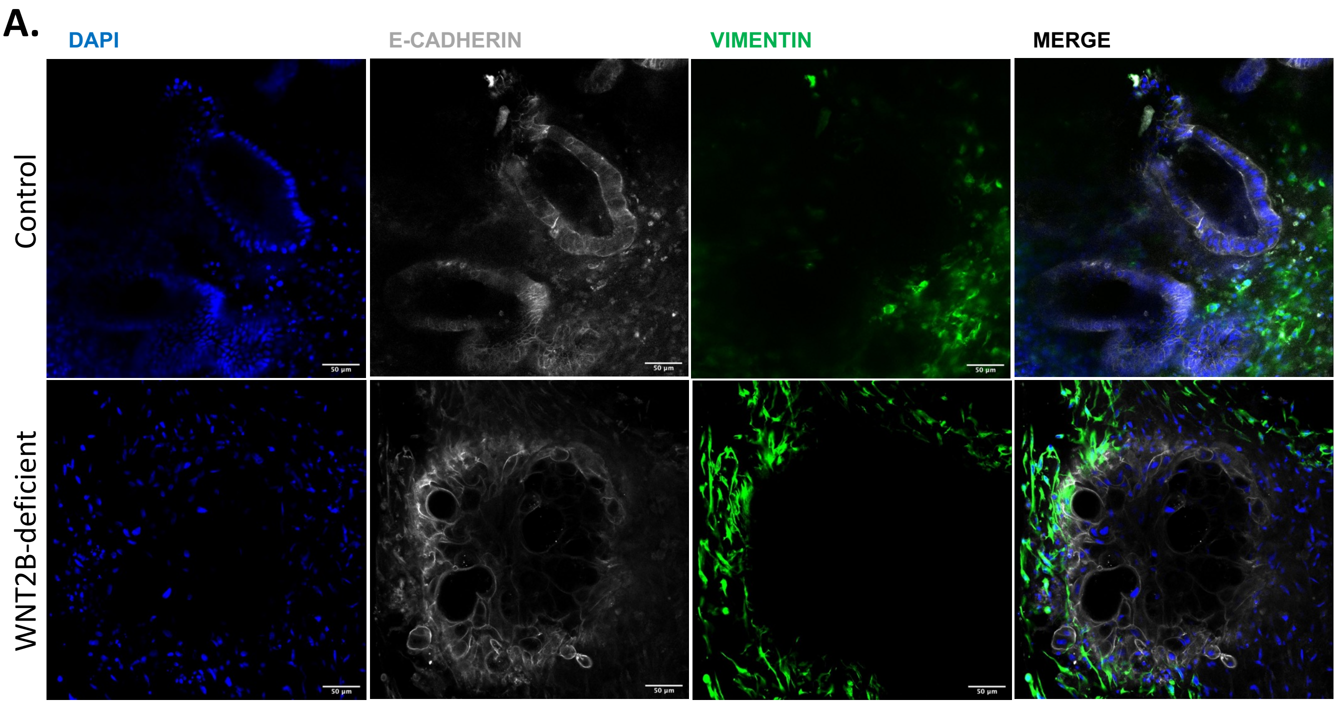


**Supplemental Figure 5**. Confocal microscopy of WNT2B-deficient HIOs. A: Expression of epithelial (E-cadherin, grey) and mesenchymal (vimentin, green) components. Scale bar = 50 microns. Scale bar = 100 microns.

**A.**

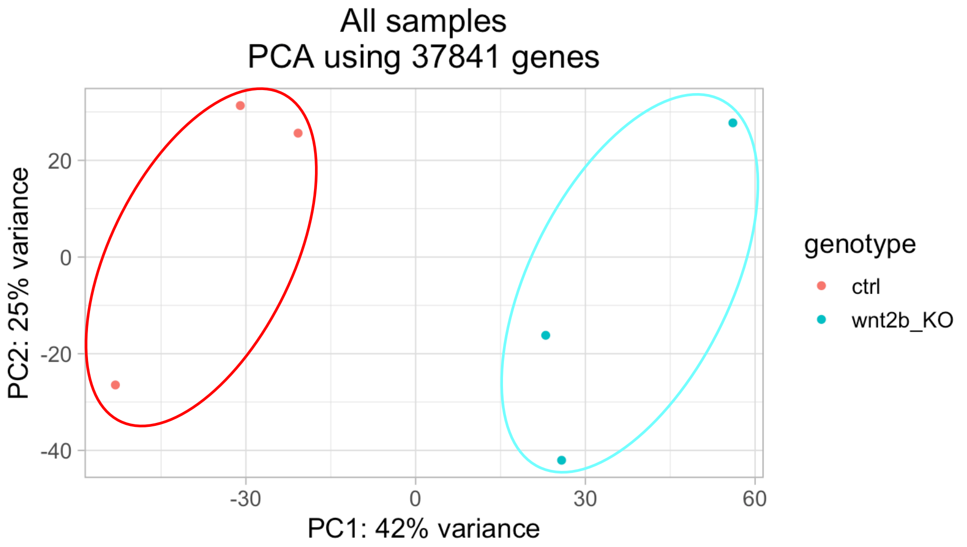


**B.**

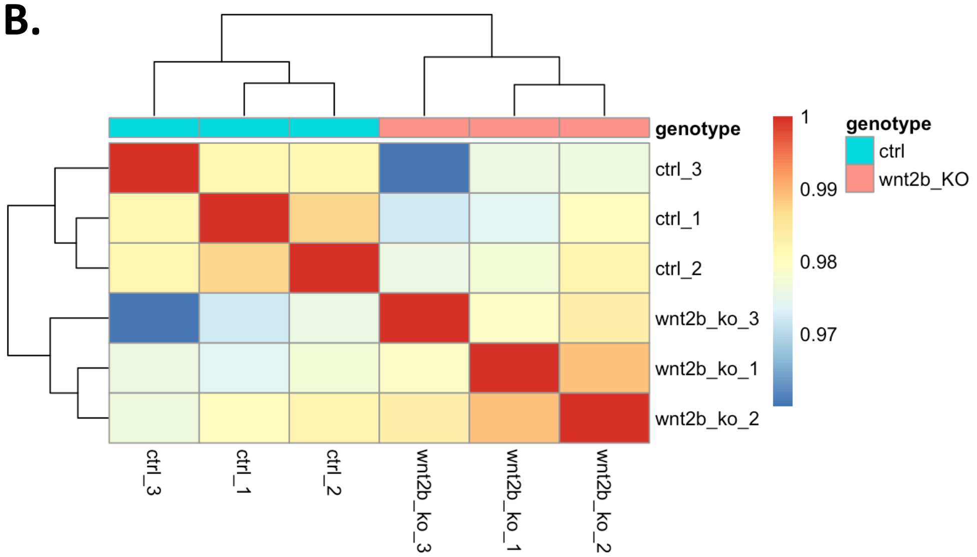


**Supplemental Figure 6**. **RNASeq quality analysis**. A. Principal component analysis for HIO RNA samples analyzed by RNASeq. B. Gene expression heatmap for hierarchical clustering of all the samples analyzed.

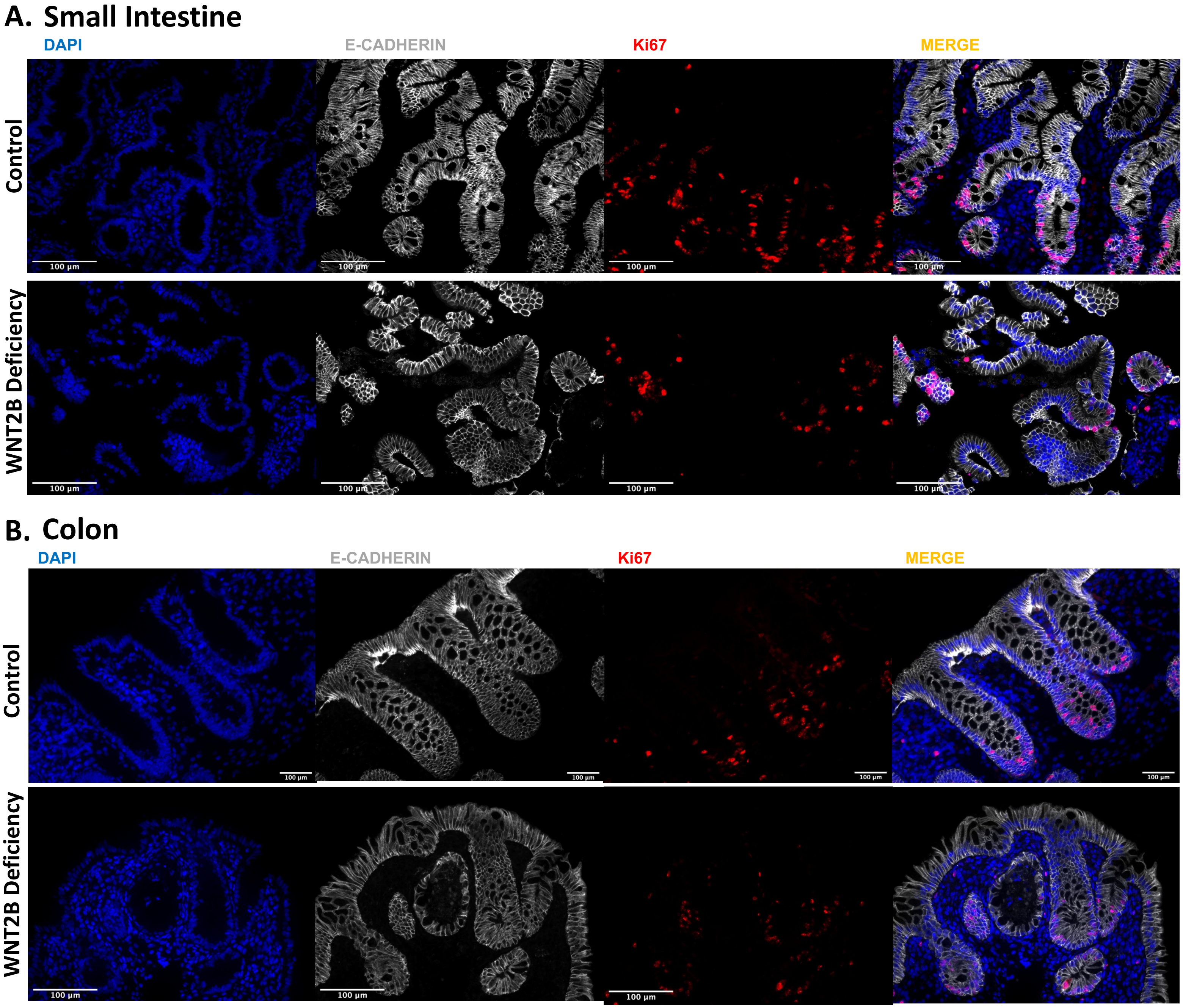


**Supplemental Figure 7**. Immunofluorescence microscopy of E-cadherin and Ki67 staining from small intestinal (A) and colonic (B) biopsies of human subject with WNT2B deficiency (hW2bD). Size bar = 100µm.
