## Supplemental Table 1 for "WNT2B Deficiency Causes Increased Susceptibility to Colitis in Mice and Impairs Intestinal Epithelial Development in Humans"

**Supplemental Table 1. Reagents**

**Primers**

| Mouse | 18S | Mm03928990_g1 | | Thermo |
| --- | --- | --- | --- | --- |
|  | Wnt2b | Mm00437330_m1 | | Thermo |
|  | LGr5 | Mm00438890_m1 | | Thermo |
|  | Lyz1 | Mm00657323_m1 | | Thermo |
|  | Muc2 | Mm00458296_g1 | | Thermo |
|  | Alpi | Mm01285814_g1 | | Thermo |
|  | Rspo3 | Mm01188251_m1 | | Thermo |
|  | Rspo1 | Mm00507077_m1 | | Thermo |
|  | Wnt3 | Mm00437336_m1 | | Thermo |
|  | Wnt5a | Mm00437347_m1 |  | Thermo |
|  | Lef1 | Mm00550265_m1 | | Thermo |
|  | Tcf4 | Mm00443210_m1 | | Thermo |
|  | Ascl2 | Mm01268891_g1 |  | Thermo |
|  | Atoh1 | Mm00476035_s1 | | Thermo |

**Antibodies**

| **Mouse** |  |  |
| --- | --- | --- |
| Ki67 | Recombinant Anti-Ki67 antibody [SP6] (ab16667) | abcam |
| **Human** |  |  |
| Ki67 | Recombinant Anti-Ki67 antibody [SP6] (ab16667) | abcam |
| E-cadherin | E-Cadherin (24E10) Rabbit mAb #3195 | Cell Signaling Technology |
| Vimentin | Recombinant Anti-Vimentin antibody [EPR3776] - Cytoskeleton Marker (ab92547) | abcam |
