## Supplemental Table 2 for "WNT2B Deficiency Causes Increased Susceptibility to Colitis in Mice and Impairs Intestinal Epithelial Development in Humans"

**Supplemental Table 2, Epithelial Organoid Culture Media**

**Murine Small Intestine (ENR) Media (100mL)**

| **Component** | **Volume** | **Catalog #** | **Final Concentration** | **Purpose** |
| --- | --- | --- | --- | --- |
| Rspondin-1 Conditioned Media | 10mL | HDDC Organoid Core | 10% | Contributes Rspondin-1 for stem cell proliferation |
| Advanced DMEM/F-12 | 85mL | Life Technologies 12634028 | 85% | Contributes glucose, NEAAs, sodium pyruvate, phenol red |
| Glutamax (100X) | 1mL | Life Technologies 35050061 | 1X | Contributes L-glutamine to support growth |
| HEPES (1M) | 1mL | Life Technologies 15630080 | 10mM | pH buffer for CO2 changes |
| Primocin (50mg/mL) | 200uL | Invivogen ant-pm-1 | 100ug/mL | Antibiotic |
| Normocin (50mg/mL) | 200uL | Invivogen ant-nr-1 | 100ug/mL | Antibiotic |
| B27 Supplement (50X) | 1mL | Life Technologies 12587010 | 0.5X | Growth supplement |
| N2 Supplement (100X) | 500uL | Life Technologies 17502001 | 0.5X | Growth supplement |
| N-Acetyl-Cysteine (500mM) | 250uL | Sigma A7250 | 1.25mM | Antioxidant |
| Noggin (100ug/mL) | 100uL | Peprotech 250-38 | 100ng/mL | BMP inhibitor |
| EGF(500ug/mL) | 10uL | Peprotech 315-09 | 50ng/mL | Epidermal growth factor, stimulates mitosis |

**Murine Colon (ENRW) Media (100mL)**

| **Component** | **Volume** | **Catalog #** | **Final Concentration** | **Purpose** |
| --- | --- | --- | --- | --- |
| NRW Conditioned Media | 50mL | HDDC Organoid Core | 50% | Contributes Noggin, Rspondin-1, and Wnt3a for stem cell proliferation |
| Advanced DMEM/F-12 | 45mL | Life Technologies 12634028 | 45% | Contributes glucose, NEAAs, sodium pyruvate, phenol red |
| Glutamax (100X) | 1mL | Life Technologies 35050061 | 1X | Contributes L-glutamine to support growth |
| HEPES (1M) | 1mL | Life Technologies 15630080 | 10mM | pH buffer for CO2 changes |
| Primocin (50mg/mL) | 200uL | Invivogen ant-pm-1 | 100ug/mL | Antibiotic |
| Normocin (50mg/mL) | 200uL | Invivogen ant-nr-1 | 100ug/mL | Antibiotic |
| B27 Supplement (50X) | 1mL | Life Technologies 12587010 | 0.5X | Growth supplement |
| N2 Supplement (100X) | 500uL | Life Technologies 17502001 | 0.5X | Growth supplement |
| N-Acetyl-Cysteine (500mM) | 250uL | Sigma A7250 | 1.25mM | Antioxidant |
| EGF(500ug/mL) | 10uL | Peprotech 315-09 | 50ng/mL | Epidermal growth factor, stimulates mitosis |

**Human NRW Organoid Media, 100 mL**

| 50 ml | NRW CM (Stappenbeck Conditioned Medium, from organoid core) |
| --- | --- |
| 30 ml | Advanced DMEM F12 (Gibco) |
| 1 ml | Glutamax (Gibco) |
| 1 ml | Hepes (Gibco) |
| 200 ul | Primocin (or Pen Strep) (Invivogen) |
| 200 uL | Normicin (Invivogen) |
| 1 ml | B27 (Gibco) |
| 0.5 ml | N2 (Gibco) |
| 1 ml | Nicotinamide (1 M) |
| 100 ul | N-acetyl cysteine (500 mM) |
| 10 ul | EGF 10000x (500 ug/ml) |
| 10 ul | Gastrin 10000x (500 uM) |
| 100 ul | A-83-01 (0.5 mM) |
| 33.2 ul | SB2002190 (5 mg in 505 ul DMSO) |
| 1 uL | Prostaglandin E2 (100 mM) |
| 100 ul | Rho kin inhibit (optional) |
