## Supplemental Table 3 for "WNT2B Deficiency Causes Increased Susceptibility to Colitis in Mice and Impairs Intestinal Epithelial Development in Humans"

Supplemental Table 3. Colitis histology scoring.

| **Score** | **Inflammatory Infiltrate** | **Goblet Cell Loss** | **Crypt Loss** | **Crypt Hyperplasia** | **Muscle Thickening** | **Submucosal Inflammation** |
| --- | --- | --- | --- | --- | --- | --- |
| 0 | none | none | normal | none | none | none |
| 1 | increased | <10% | decreased by <10% | slight increase | slight | individual cells |
| 2 | also in submucosa | 10-50% | decreased 20-50% | 2-3x increase | strong | infiltrates |
| 3 | transmural | >50% | decreased >50% | >3x increase | excessive | large infiltrates |
