## Supplemental Methods for "WNT2B Deficiency Causes Increased Susceptibility to Colitis in Mice and Impairs Intestinal Epithelial Development in Humans"

*Rigor and Reproducibility*. Experimental data for mouse experiments is representative of a minimum of three experiments with similar outcomes. Experimental data for HIO experiments is also representative of two replicates. Staining from the human patient biopsies represent two biological replicates.

*Murine Organoid Generation.* Intestinal tissue, either duodenal or colonic, was cleaned using cold intraluminal phosphate buffered saline (PBS), cut into smaller pieces, and cleaned in PBS until the supernatant was clear. The tissue was then placed in 2mM EDTA (small intestine) or 5mM EDTA (colon) and incubated on ice with rocking for 30 minutes. The tissue was then shaken vigorously for 2 minutes to release crypts and pipetted up and down 25 times with a 10mL serological pipet while mixing. The solution was filtered and centrifuged. Dissociated cells were then washed in Advanced DMEM/F12 (Gibco) and reconstituted in Matrigel (Corning, 50μL/well) plated in 24-well plates and incubated for 10 minutes at 37ºC. Next, 0.5mL growth media was added to the wells (Supplemental table 2).

*Human iPS Generation*. Fibroblasts from a 5 year old male donor with Wnt2b deficiency [(hg19) chr 1:113057518 (GenBank: NM_024494; c.205C>T [p.Arg69*])] were cultured in MEF media (DMEM, 10% FBS, 1mM non-essential amino acids). Cells at ~50% confluency were transduced overnight with Sendai viral vectors (Cytotune 2.0, ThermoFisher Scientific) at MOIs of 2.5 (Klf4,Oct4,Sox2), 2.5 (cMyc), and 1.5 (Klf4).  Spent media was removed from cells and completely replaced with fresh MEF media on days 1,3, and 5 post-transduction. On day 7, transduced cells were plated in MEF media on irradiated MEF feeders (187,500 cells/well) in 6 well plates coated with 0.1% gelatin. On day 8, spent MEF media was removed and replaced with hESC media (DMEM/F12, 20% knockout Serum replacement, 1mM L-Glutamine, 0.1mM beta-mercaptoethanol, 1x non-essential amino acids, 2µg/mL bFGF). Starting on d8, wells underwent a complete daily media change with 2.5mL hESC media. Putative iPSC colonies were then manually excised and replated in feeder free culture conditions consisting of Stem cell qualified Cultrex (BioTechne) and mTeSR1 (StemCell Technologies). Lines exhibiting robust proliferation and maintenance of stereotypical human pluripotent stem cell morphology were then expanded and cryopreserved at ~ passage 10.

*Human Intestinal Organoid Cultures.* The hiPSC colonies were gently dissociated into single cells with Accutase (StemCell Technologies, 07920) and seeded on Matrigel coated 24-well plates at a density of 30-60K per well. The cells were cultured in mTeSR1 medium with 10uM ROCK inhibitor Y27632 (Tocris, 1254) for 24 hours, then replaced with fresh mTeSR1 medium. After 24 hours, the cells were treated with 100 ng/mL recombinant human Activin A (R&D Systems, 338-AC-050) in RPMI 1640 media (Gibco) supplemented with 2mM L-Glutamine (Gibco) and fetal bovine serum (Gibco, day 1: 0%, day 2: 0.2% and day 3: 2%) for derivation into definitive endodermal cells. After differentiation into endodermal cells, the cells were treated with 500 ng/mL recombinant human FGF4 (R&D Systems) and 3μM ChiR 99021 (R&D Systems, 4423) in RPMI 1640 media supplemented with 100 U/mL Penicillin-Streptomycin (Gibco, 15140122), 2mM L-Glutamine (Gibco, A2916801), and 2% of fetal bovine serum for 3 days for derivation into midgut/hindgut cells. On day 3, the cells were gently dissociated into single cells with Accutase and seeded on pretreated Aggrewell plates (StemCell Technologies, 34450) at a density of 3.6 x 10^6 cells per well. The cells were cultured in midgut/hindgut differentiated media for 24 hours. Then three-dimensional spheroids were collected and embedded in Matrigel on 24-well culture plates. After the Matrigel had solidified, advanced DMEM/F12 (Gibco, 12634-010) supplemented with 2 mM Glutamax (Gibco, 35050-061), 10 mM HEPES (Gibco, 15630080), 2% (v/v) B27 supplement (Gibco, 17504044), 1% (v/v) N2 supplement (Gibco, 17502048), 100 U/mL Penicillin-Streptomycin, and 100 ng/mL recombinant human EGF (R&D Systems, 236-EG, only on the first 3 days), and the growth factors described above were overlaid and replaced every 3 days for more than 28 days for derivation into intestinal lineages.

*Immunofluorescence.* Slides were prepared for staining by immersing in Trilogy buffer and placing in a pressure cooker for 5 minutes. Slides were then rinsed with Trilogy buffer and washed with deionized water and then rinsed further in PBS. Blocking buffer was added to each slide (Blocking buffer: 0.3% Tween-20; 10% normal donkey serum; 0.05% BSA in PBS) and incubated at room temperature for 1 hour, gently shaking. Primary antibody was then added in 1% NGS-PBS at 4°C overnight according to the concentration in Supplementary Table 1. Slides were then washed in PBS then incubated with the secondary antibody also per Supp. Table 1 for 60 minutes at 37°C. DAPI 1:1000 was added for 10 mins and then the slides were washed again and sealed with a coverslip.
